# Multiplex shRNA Screening of Germ Cell Development by *in vivo* Transfection of Mouse Testis

**DOI:** 10.1101/046391

**Authors:** Nicholas R. Y. Ho, Abul R. Usmani, Yan Yin, Liang Ma, Donald F. Conrad

## Abstract

Spermatozoa are one of the few mammalian cells types that cannot be fully derived *in vitro*, severely limiting the application of modern genomic techniques to study germ cell biology. The current gold standard approach of characterizing single gene knockout mice is slow as generation of each mutant line can take 6-9 months. Here, we describe an *in vivo* approach to rapid functional screening of germline genes based on a new non-surgical, non-viral *in vivo* transfection method to deliver nucleic acids into testicular germ cells. By coupling multiplex transfection of short hairpin RNA constructs with pooled amplicon sequencing as a readout, we were able to screen many genes for spermatogenesis function in a quick and inexpensive experiment. We transfected nine mouse testes with a pilot pool of RNAi against well-characterized genes to show that this system is highly reproducible and accurate. With a false negative rate of 18% and a false positive rate of 12%, this method has similar performance as other RNAi screens in the well-described *Drosophila* model system. In a separate experiment, we screened 26 uncharacterized genes computationally predicted to be essential for spermatogenesis and found numerous candidates for follow up studies. Further, by characterizing the effect of both libraries on neuronal N2a cells, we show that our screening results from testis are tissue-specific. Our calculations indicate that the current implementation of this approach could be used to screen thousands of protein-coding genes simultaneously in a single mouse testis. The experimental protocols and analysis scripts provided will enable other groups to use this procedure to study diverse aspects of germ cell biology ranging from epigenetics to cell physiology. This approach also has great promise as an applied tool for validating diagnoses made from medical genome sequencing, or designing synthetic biological sequences that act as potent and highly specific male contraceptives.

## Introduction

As the carrier of genetic information from one generation to the next, germ cells are derived from one of the most specialized developmental processes in the body. Once the stem cell commits to the terminal sperm lineage, it has to trigger various transcription factors to begin specialized metabolic processes, produce haploid cells via meiosis, tightly pack the genomic DNA and prepare for functions like flagellum motion, cell recognition and acrosome formation. Furthermore, some of the pathways for core physiological processes are distinct from somatic cells despite having similar functions (Eddy 1998). Perhaps due to this richness, the study of mammalian spermatogenesis has led to numerous seminal discoveries with broad implications in areas of biology such as stem cells (Chen *et al*. 2005), transposable elements (Girard *et al*. 2006), adaptive evolution (Carelli *et al*. 2016) and speciation (Good *et al*. 2010).

Despite the opportunities for discovery in the field of spermatogenesis, the pace of progress has been limited because existing *in vitro* model systems are technically challenging to implement (Stukenborg *et al*. 2009; Sato *et al*. 2011; Dores and Dobrinski 2014). Generation of knockout mouse models has thus been the most popular tool to characterize the function of genes in germ cells. Due to the high cost (over $5,000 USD) and the time involved (over 1 year) in deriving a colony of a new mouse line, the “one-gene, one-mouse” approach cannot be easily used to perform systematic screens of the genome. This limited access to high-throughput screening in germ cells is a stark contrast to the rapid expansion of multiplex genomic techniques now being used in cell lines (Consortium 2012). As these large-scale, multiplex genomic studies become more commonplace, the gap between our knowledge of germ cell and somatic cell biology will only grow if single-mutation mouse models remain the method of choice.

To address this problem, we have developed a quick, simple, and inexpensive method to screen and verify numerous genes simultaneously *in vivo* for spermatogenesis function. The mammalian testis continuously produces millions of mature sperm each day. This abundance of testicular germ cells would easily support a multiplex genomics screen like those used in cell lines if one could develop a viable way to deliver nucleic acids into the testis of a living animal. The basis for our approach is a novel method for direct transfection of testicular germ cells, coupled to the popular RNA interference (RNAi) screen, a mature technology commonly used to efficiently elucidate gene function. RNAi screens have been used in cell lines (Luo *et al*. 2008; Zuber *et al*. 2011b) or *in vivo* (Zender *et al*. 2008; Bric *et al*. 2009; Meacham *et al*. 2009; Zuber *et al*. 2011a; Beronja *et al*. 2013; Wuestefeld *et al*. 2013) in somatic tissues to discover important genes for a variety of biological processes.

Here we demonstrate the feasibility of using this low cost transfection method in mouse testes to screen multiple genes simultaneously for functional importance in spermatogenesis. By carefully designing the pilot study, we were also able to benchmark this system to prove the importance of large numbers of biological replicates and quantify the limits of this system. We also applied this method to establish the functional importance of twenty six uncharacterized genes that we previously predicted to be important for infertility via machine learning (Ho *et al*. 2015).

## Results

### In vivo transfection

Recently, a small number of studies have shown that one can use viruses to transfect mouse tissues *in vivo* with a sufficiently high transfection rate for multiplex selection experiments (Beronja *et al*. 2013; Wuestefeld *et al*. 2013). One of us (A.U.) recently developed a novel way for direct transfection of testicular germ cells that provides a non-viral, non-surgical alternative for nucleic acid delivery. This technique uses a buffered salt solution to generate an osmotic gradient that drives water and dissolved linear DNA into the germ cells of the testis at a reasonably high rate (Usmani *et al*. 2016). Once inside the cells, the DNA integrates spontaneously and randomly into the genome at a concentration-dependent rate, presumably through non-homologous end joining during routine DNA repair. We adapted this approach to transfect the germ cells in the testis to render it compatible with linear DNA libraries expressing small hairpin RNAs (shRNAs) **(Methods, Figure 1)**. Visual inspection of testis transfected with a GFP reporter indicated that both somatic and germ cell types receive construct, but that germ cells, which comprise well over 75% of the adult testis, are preferentially transfected **(Figure 2)**. By integrating the expression cassette into the genome, we achieved stable expression of the RNAi construct in transfected cells that could persist for weeks and could be observed in germ cells spanning all stages of spermatogenesis (Figure 2).

**Figure 1.**
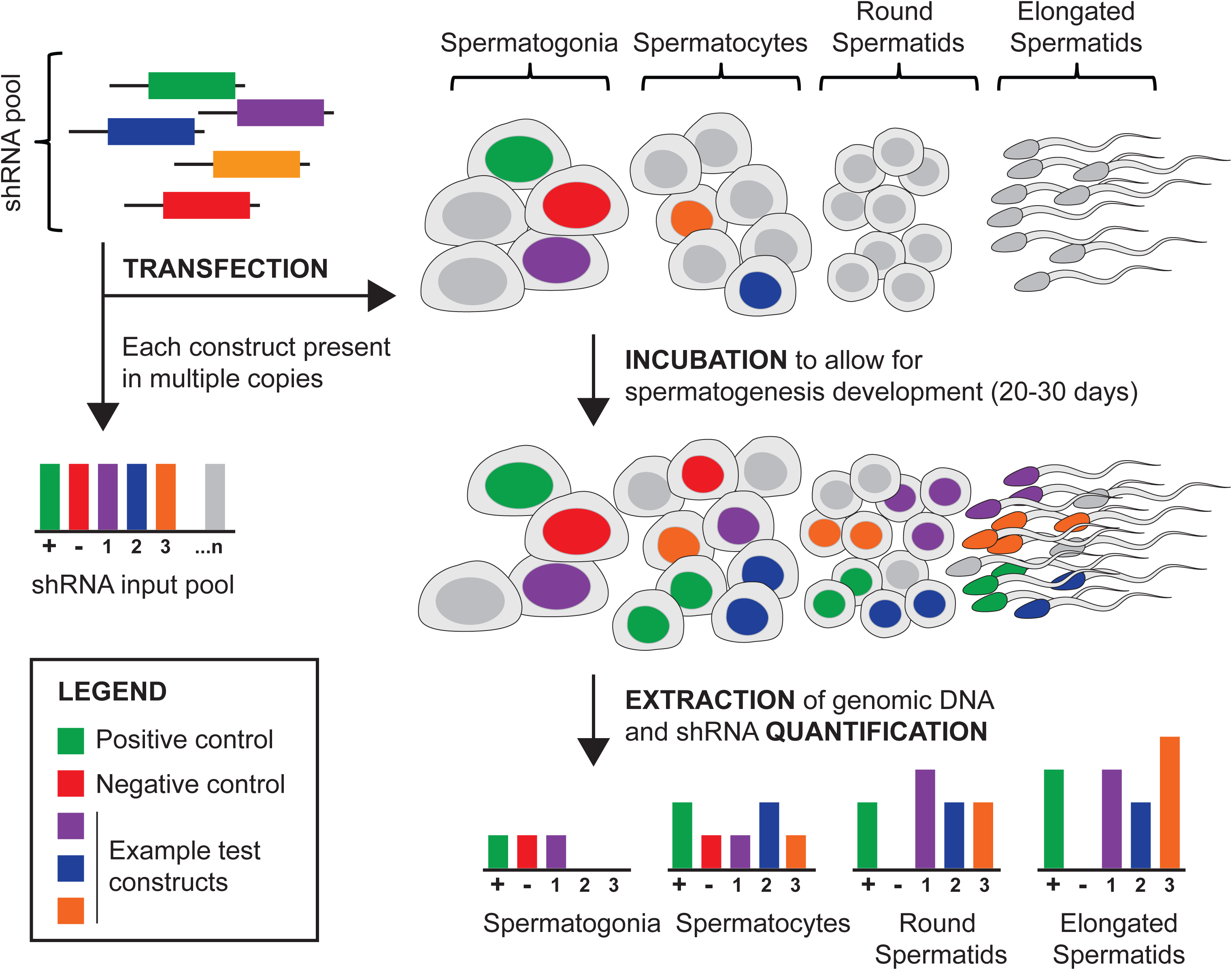
**Overview of Experimental approach**. An shRNA library is designed against 20-30 genes, comprising 3-5 distinct constructs per gene. For a typical library, 3 types of genes are selected-positive controls (genes known to be essential for spermatogenesis), negative controls (genes not thought to have a testis phenotype when biallelically inactivated), and “test” genes whose role in spermatogenesis is uncertain and to be determined in the screen. This library is maintained as a pool of plasmids, each plasmid bearing one shRNA construct. DNA from the input pool is isolated and split into two aliquots: one for quantification and one for transfection. The transfection aliquot is delivered to the testis using our non-surgical approach (Methods). Twenty to thirty days following transfection, the DNA from germ cells are isolated and shRNA-specific primers are used to create sequencing libraries from both the input pool and the post-testis pools. Sequencing libraries are then sequenced using short-read sequencing and the relative abundance of each shRNA construct in a library can be estimated by counting the sequencing reads containing that shRNA construct. The effect of a gene knockdown on spermatogenesis is summarized as the log2 fold change of shRNA abundance in the post-testis pool compared to the input pool. The screen is designed to be used in combination with any cell selection scheme; for instance libraries can be made from purified subpopulations of germ cells to identify stage-specific effects. In the present study, we largely report results from post-testis libraries made from total testis DNA.

**Figure 2.**
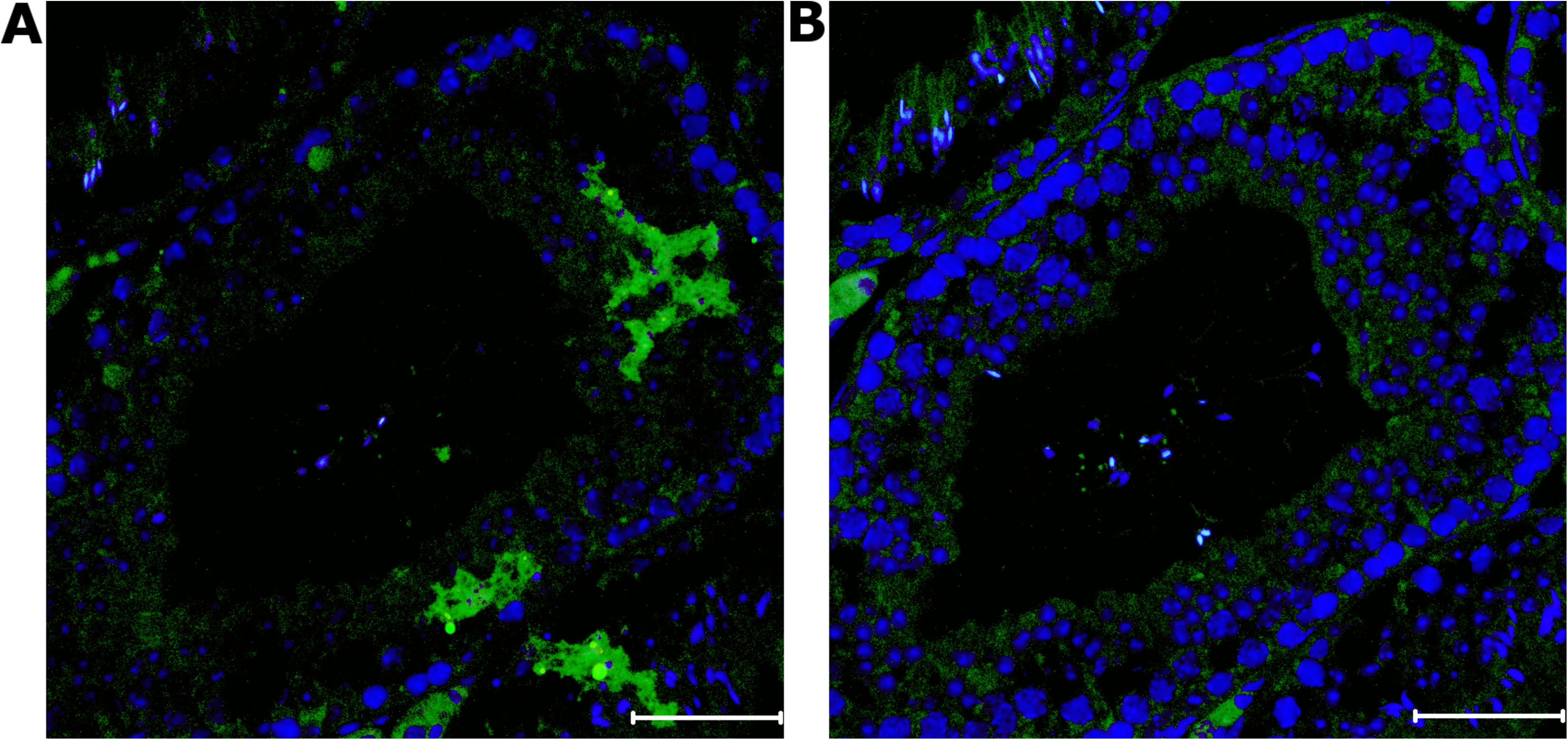
**Expression of a transgene delivered directly to testicular germ cells with TrisHCl transfection. (A)** a C57Bl/6J mouse testis injected three times with 15µg of a plasmid constitutively expressing GFP via a CMV promoter dissolved in 150mM TrisHCl with 3-4 days between each injection. Using an anti-GFP antibody, GFP was observed in numerous germ cell subtypes including spermatids, spermatocytes and spermatogonia. **(B)** an adjacent section of the same testis, processed without primary antibody against GFP. The scale bar on the bottom right indicates 50µM in length. Blue indicates Hoechst 33342 and green indicates GFP.

### Pilot shRNA screen

In order to optimize the screening technique, we designed a pilot shRNA pool targeting 4 classes of genes: i) sixteen genes that have been shown to cause assorted dosage-dependent defects of sperm development when inactivated and/or overexpressed (BAX, CSF, KIT, PIN1, CPEB1, GNPAT, MLH3, SPO11, CIB1, MAP7, PYGO2, TBPL1, SH2B1, TSN, SIRT1, VAMP7 & VDAC3); ii) five genes that have been characterized with knockout mice but have not been linked to spermatogenesis defects (MMP3, SYT4, TFF3, TNFSF4, TYRP1); and iii) three genes that have not been characterized in a mouse model and are not expressed in mouse testis (APOC4, LCE1I, SCRG1) Our pool contained 119 unique RNAi constructs with a mode of five RNAi constructs per targeted gene **(see Table S1)**.

Using the pilot pool, we first benchmarked the infection efficiency of our novel osmotic gradient transfection buffer (“Tris-HCl + DNA”) against the current gold standard for *in vivo* screening, lentiviral infection. To enable rapid measurement of infection efficiency, we designed a qPCR assay for DNA copy number of the shRNA constructs, calibrated to actin (see **Figure S1**). We compared the infection rate of a single injection of high-titer virus (109 Tu/ml) against either a single injection of Tris-HCl with 15µg DNA, or a single injection of Tris-HCl + DNA followed by 4 “booster” shots of additional 15µg aliquots of shRNA library separated 3-4 days apart (**Methods**). A single injection of high titer lentivirus produced a testis with a higher infection rate than a single injection of Tris-HCl with 15µg of DNA **(see Table S2)**. However, the single injection of lentivirus produced an infection rate was lower than five injections of the Tris-HCl + DNA (0.1% versus 1%-4%). Due to the lower infection rate, we observed low reproducibility and more dropout of shRNA constructs in the single DNA and viral injection testes samples compared to the five DNA injection testes. Since the cost of performing five DNA injections is significantly lower than even one high-titer lentivirus injection (about $50 versus $250), we decided to proceed with the direct DNA injections rather than attempt multiple lentiviral injections.

Next, we transfected the pilot library into 9 mouse testes using the 5-shot protocol in order to perform a screen for genes required for germ cell differentiation. After transfection we waited 20 days, slightly longer than half of one cycle of spermatogenesis, before recovery and quantification of the shRNA library from the testis. We extracted total genomic DNA from each testis, and used PCR to create at least 3 replicate sequencing libraries from each testis **(Methods, Figure S2)**. The frequency estimates of individual shRNAs were very strongly correlated among biological replicates (Pearson correlations 0.66-0.96, **Figure 3**).

**Figure 3.**
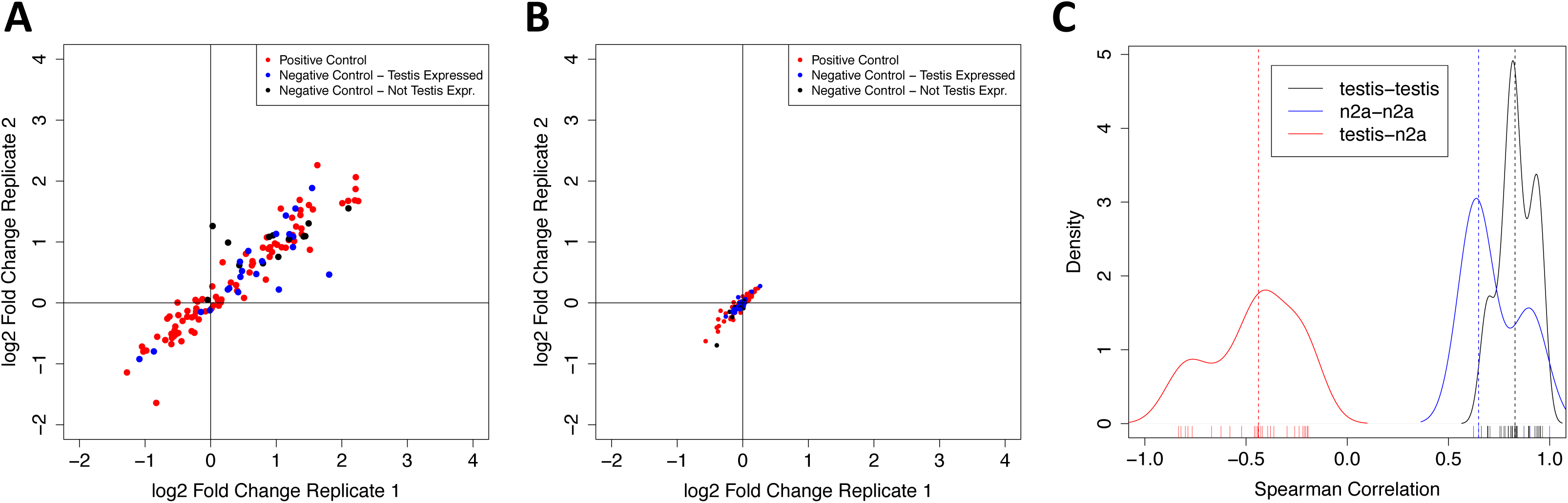
**shRNA screening assays are highly reproducible and tissue-specific**. We performed numerous biological replicates of the pilot library screening on testis (n=9) and the n2a neuronal cell line (n=3). **(A)** Comparison of two biological replicates within the testis. For each replicate, we summarize the frequency of all 119 shRNA constructs as the log2 ratio of testis frequency/input library frequency. Overall, the concordance of fold-changes between these two replicates is high (Spearman R=0.94). While 11% of negative control constructs show negative fold changes in both biological replicates, all of these correspond to genes expressed in testis. On the other hand, 35% of positive control constructs show negative fold changes. **(B)** Comparison of two biological replicates of the screen in n2a cells shows the screen is also high reproducible in this cell population (Spearman R=0.90). The variance in fold change among constructs is visibly smaller in the n2a experiments when compared to the testis; this is likely due to higher transfection efficiency in n2a cells. **(C)** As a broader summary of screen reproducibility, we calculated Spearman correlations between all pairs of biological replicates for testis (black line) and n2a cells (blue line), as well as all possible pairs of n2a and testis replicates (red). Biological replicates of the same source tissue were highly correlated (median Spearman R for n2a-n2a comparisons= 0.65, dashed blue line; testis-testis = 0.83, dashed black line). However, screen results from different sources were moderately to strongly anticorrelated (median Spearman R=-0.44, dashed red line). This anticorrelation was largely driven by “negative control” shRNAs showing depletion in n2a cells, as well as neutral behavior of positive controls in n2a cells.

We devised an analysis strategy to identify shRNAs that were significantly depleted in the post-testis shRNA library when compared to the input library (**Methods**). We used a p-vaue of 0.01 to declare any given construct as depleted in the post-testis library. In order to reduce the chance of off-target effects, we required any given gene to have two different significantly depleted shRNAs before declaring that gene as essential for spermatogenesis. Given the low multiplicity of infection of our transfection system, the fold-changes of each shRNA construct should be independent random variables under the null hypothesis that there is no relationship between shRNA expression and germ cell survival. Thus, the probability of observing a gene with 2 depleted shRNA constructs should be 1 x 10-4 or (0.01)2.

Next, we set out to estimate the false negative and false-positive rate of our shRNA screening system when using this “two-hit” criterion. Of 17 positive control genes, 14 were identified as required for spermatogenesis, producing an estimate of 18% for the false negative rate of our approach **(Figure 4A)**. The positive controls that did not show significant depletion were BAX, MATP7 and SH2B1. These false negatives could not be simply attributed to low sequence coverage, as the average coverage of constructs for each gene was 6188X, 1732X and 1163X respectively. Five out of 5 (100%) of BAX constructs were experimentally validated to knock down expression; likewise 4/5 of the MTAP7 and 5/5 of the SH2B1 constructs were experimentally validated. One of the eight negative control genes, TNFSF4 (which is moderately expressed in the testis), was also significantly depleted, producing a conservative false positive rate estimate of 12.5% (1/8).

**Figure 4.**
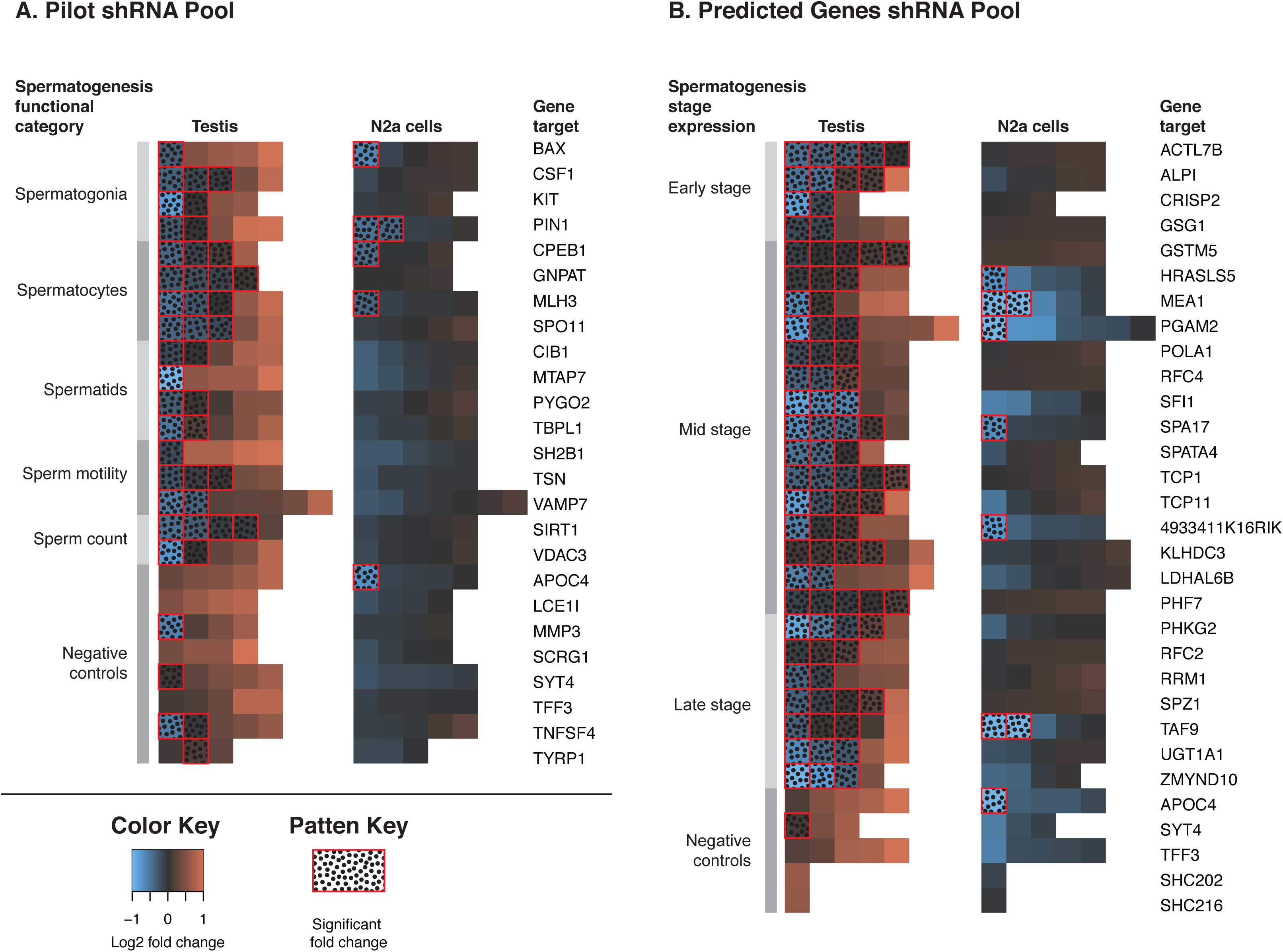
**shRNA screen results** Overview of the full screening results for (A) the pilot shRNA pool and (B) the predicted genes shRNA pool. Each row of the heatmap summarizes the screening data for constructs targeting a single gene that were transfected into either testis (left heatmap) or n2A cells (right heatmap). Each cell shows the log2 fold change of a single shRNA construct when comparing frequencies from the transfected tissue to the input pool frequencies. Constructs with significant depletion in testis or n2a cells are encased in a red box and further shaded by a “dot” pattern. The constructs within each row are sorted left-to-right in ascending order based on the log2 fold change. In (A), the genes are further organized into functional categories as indicated by the greyscale bar to the left of the heatmaps, and in (B) genes are also organized based on cell type of peak expression level, with stages encoded by the greyscale bar.

One drawback of our screening strategy is that the genes we identify could be required specifically in spermatogenesis, or just for general survival of all cells. We reasoned that it should be possible to disentangle these two effects by transfecting the same shRNA pool into an unrelated cell line and performing a survival screen on the cells. Genes required for general cell survival will be identified by both screens, while genes with germ-cell specific function will not.

We transfected three separate batches of Neuro-2a (N2a) cells, prepared sequencing libraries, and analyzed the data through the same pipeline as the testis samples **(Methods)**. As with the testis libraries, there was robust correlation in shRNA fold-changes among biological replicates **(Figure 3B)**. Interestingly, the shRNA fold-changes in the n2a library were moderately anti-correlated with the testis library fold-changes, indicating that we do indeed observe tissue-specific selection in our screen **(Figure 3C)**. This anticorrelation was largely driven by “negative control” shRNAs showing depletion in n2a cells, as well as neutral behavior of positive controls in n2a cells. Furthermore none of the shRNAs that were significantly depleted in the testes samples were also significantly depleted in the cell line **(Figure 4A)**. We thus concluded that the genes with multiple shRNAs depleted in testes are essential for germ cell development and not just general cell viability.

### Screening for uncharacterized predicted fertility genes

We previously developed a computational algorithm to predict genes required for spermatogenesis (Ho *et al*. 2015). For each gene in the mouse genome, this algorithm uses high-throughput genomic data to reports a probability that the gene is required for spermatogenesis. While we have benchmarked the performance of this method by comparison with genes known to be irrelevant or essential for sperm production, we never experimentally validated any of our novel predictions. We designed a new shRNA pool against the top 26 genes from our computational screen that had no known role in spermatogenesis **(Methods)**. The new shRNA pool comprised 130 shRNAs against the candidate genes and 15 negative control shRNAs. These candidates display a variety of expression signatures in spermatogenesis: four genes begin expression early in spermatogenesis (ALPI, POLA1, RFC1, RRM1), fifteen genes begin expression in the middle of spermatogenesis (CRISP2, GSTM5, HRASLS5, KLHDC3, LDHAL6B, PGAM2, PHF7, PHKG2, RFC2, SFI1, SPATA4, TAF9, TCP1, TCP11, ZMYND10), and seven genes are only expressed late in spermatogenesis (4933411K16Rik, ACTL7B, GSG1, MEA1, SPA17, SPZ1, UGT1A1) **(see Table S1)**.

Similar to the pilot screen, the standard error in shRNA frequencies estimated from technical replicates indicated strong reproducibility (Pearson correlations 0.54-0.96) and low experimental noise. None of the 15 negative control shRNAs were significantly depleted (P ≤ 0.01) while all of the 26 candidate genes had at least two shRNAs targeting them significantly depleted in the testes samples. A few genes even had up to four different significantly depleted shRNAs (P ≤ 0.01) **(Figure 4B)**. Similar to the pilot pool, we also transfected this pool into N2a cells to eliminate the possibility that these genes are required for general cell survival. Overall, the normalized read counts of the shRNA pool in N2a cells were not well correlated with the testis read counts **(Figure 4B)**. Five genes (HRASLS5, MEA1, PGAM2, SFI1 and TAF9) had two or more shRNAs that were significantly depleted (P ≤ 0.01). Given this, we concluded that most of the predicted genes (21/26) affect spermatogenesis and not survival.

### Design of future screens

Given the strong results obtained with our first two shRNA screening experiments, we believe there is great potential for this approach moving forward. In order to improve the design of future screening experiments, we reanalyzed all of the sequencing data generated from our pilot screen to characterize the relationship between both (a) the number of biological replicates performed, (b) the number of constructs screened and the power to detect significant depletion of shRNAs against a typical positive control gene. To address (a), we estimated the fraction of positive control genes detected as significantly depleted as function of number of biological replicates. To address (b) we estimated the fraction of positive control genes detected as significantly depleted as a function of sequencing coverage for each library. As in our initial analyses, we required that two shRNAs for the same gene show significantly depleted in order to declare a gene as a “hit” in the screen.

Increasing the number of biological replicates provides more confidence that the RNAi fold changes are biologically significant, but we observed diminishing returns with increasing numbers of replicates. For lenient (p<0.1), standard (p<0.05) and stringent (p<0.01) p value cutoffs there was little improvement in detectable RNAi fold change beyond 3-4, 4-5, and 6-7 biological replicates respectively (Figure 5A).

**Figure 5.**
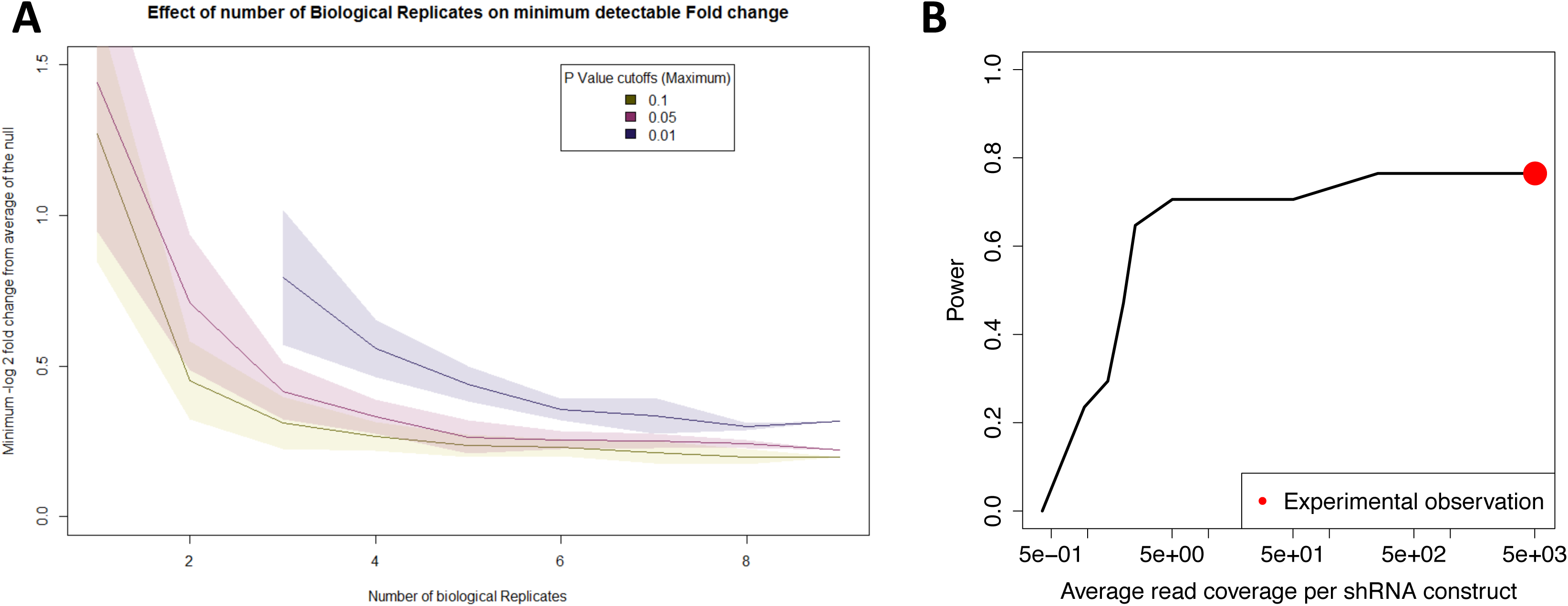
Influence of experimental design parameters on the power of the screen. We reanalyzed all of the sequencing data generated from our pilot screen to characterize the relationship between both the number of biological replicates performed, the number of constructs screened and the power to detect significant depletion of shRNAs against a typical positive control gene. (A) As the number of biological replicates increases, the minimum detectable shRNA fold change (i.e. effect size) decreases. The 3 lines indicate the median minimum log2 fold change that was declared as significant at a threshold of p ≤ 0.1 (yellow), p ≤ 0.05 (red) and p ≤ 0.01 (blue). The shaded area around each line defines a 1 standard deviation confidence interval. Note that the blue lines starts at 3 biological replicates because there were no significant observations with 2 or fewer replicates. (B) Power to detect a positive control gene as significantly depleted increases with increasing average read coverage of each shRNA construct. The average coverage and power of the full pilot experiment is indicated with a red dot.

Each shRNA construct was allocated an average of 5,000 sequencing reads in our actual experiments. When using 9 biological replicates, we found that we could have reduced the average read coverage to 250 reads per construct without any loss of power, and that 70% power could be maintained to even 5 reads per construct (Figure 5B). Reducing average coverage further rapidly and drastically reduces power.

## Discussion

The combination of multiplex genetic manipulation and high-throughput phenotyping presents incredible opportunities for studying the genome biology and evolution of the mammalian germline. Here we have presented a method that opens the door to *in vivo* genomic experiments in mammalian testis and demonstrate its utility by performing multiplex shRNA screens for genes essential to spermatogenesis. The performance of our direct *in vivo* screen is comparable to other benchmarked RNAi screens performed in other model systems. Our false negative rate of 18% is up to the standards of various studies benchmarking RNAi screens for different pathways in *Drosophila melanogaster* (*DasGupta et al*. *2007; Liu et al*. *2009; Booker et al*. *2011; Hao et al*. *2013*) where it was reported to be between 13% and 50%. Our false positive rate of 12.5% also compares favorably to the same study (Liu *et al*. 2009) which found their false positive rates to be between 7% and 18%.

As it stands the technique produces strongly reproducible results but several aspects of the system could be improved. The key limitation is transfection efficiency. *In vivo* transfection rates tend to be orders of magnitude lower than *in vitro* systems for the same transfection reagent (Li and Huang 2000). This leads to trade-offs between the size of the screening library, the number of biological replicates and the cost of the transfection reagent. To avoid the issue of low transfection rates in other tissues, some groups have transfected cells *in vitro* and then transplanted them into recipient mice and performed screening in the xenografted models (Zender *et al*. 2008; Bric *et al*. 2009; Meacham *et al*. 2009; Zuber *et al*. 2011a). In principle this may work for male germ cells, as spermatogonial stem cells have been transplanted into sterile donor testes to restore fertility in various species (Brinster and Zimmermann 1994; Ryu *et al*. 2007). In practice, this stem cell transplantation is a difficult technique to perform and a low number of unique stem cells successfully transplant per mouse. This creates a bottleneck that makes cell transplantation inappropriate for a screening study, since screening relies on having a large number of independent transfection events in order to produce statistically significant results. More technically challenging protocols such as efferent duct injection or in-vitro organogenesis could enhance transfection efficiency. It could also be possible to use experimental evolution to generate new viral serotypes with high trophism for testis.

We were extremely conservative with the multiplexity of the pool in this study. In order to avoid dropout of individual constructs, and produce reproducible signals, we ensured that the number of cells transfected was orders of magnitude larger than the number of shRNAs. In this manner we could be confident that no shRNA would be underrepresented in the final pool due to transfection efficiency. Even with a transfection rate of 1-5%, we were able to obtain consistent signals using pools consisting of up to approximately 150 shRNAs. It is estimated that there are a total of 108 cells in the adult murine testis (Tegelenbosch and de Rooij 1993). With a transfection efficiency of 1%, 106 cells should receive constructs, suggesting that in our pilot experiment, each of our 119 unique shRNA constructs infected an average of 8,403 cells. Conservatively, if we require that each construct transfect 100 independent cells, we could increase the scale of the screen to 10,000 constructs. This would allow a screen of 2,000 genes with 5 constructs per gene.

Another limitation of the present study is that our transfection approach is not able to target specific cell types; this presents challenges for interpretation of screen results and for confounds our ability to identify the function of genes with stage-specific effects on spermatogenesis. A simple way to address this limitation of “off-target” transfection would be to make shRNA constructs expressed by an inducible promoter (e.g. Cre) and transfect the library into an animal expressing the inducer in a stage-specific manner (e.g. a Vasa-Cre or Stra8-Cre animal).

As a proof-of-principle, we used our screening system to test 26 genes that we previously predicted as being essential for spermatogenesis. We validated all 26, which is remarkable, but also consistent with the fact that these were selected as the highest ranked (but unknown) spermatogenesis genes from over 1,000 possible candidates. There are diverse molecular functions represented in this set of 26. Seven genes are thought to be involved in DNA replication and chromatin dynamics (ACTL7B, KLHDC3, POLA1, RFC2, RFC4, RRM1, SFI1), seven in metabolism (ALPI, GSTM5, HRASLS5, LDHAL6B, PGAM2, PHKG2, UGT1A1), four transcription factors (OHF7, SPZ1, TAF9, ZMYND10), two protein folding (TCP1, TCP11), one cell binding in sperm (SPA17), and one involved in capacitation of the membrane (CRISP2). In order to better understand our ascertainment, we visualized the functional relationships of these candidate genes with other known spermatogenesis genes **(see Figure S3)**. Most of the candidates were related through a single large network of interactions, but, interestingly, all 7 candidates implicated in DNA replication and chromatin dynamics were related by a smaller, standalone network that has yet to be characterized in testis. Also noteworthy were the 3 genes of unknown molecular function - 4933411K16Rik, GSG1, and SPATA4. These genes could work in novel pathways, providing new insights about spermatogenesis.

Beyond its use as a validation tool, there are many potential applications of testis shRNA screening. During this project we attempted to enhance the characterization of our target genes by performing additional screens that use cell stage or functional separation (via FACS and sperm motility assays respectively). Unfortunately, because the number of cells we could retrieve in this manner was limited, we were unable to prepare sequencing libraries from the subpopulations. If we could transfect a majority (60-80%) of the cells, or if it was possible to sort and retrieve large numbers of cells (10s of millions), direct functional assays using an RNAi pool may be possible. There are many mysterious aspects of germ cell development and function that could be rapidly screened when *in vivo* transfection becomes more efficient – what are the causes and consequences of germ cell epimutation? What are the determinants of sperm morphology, and why is sperm morphology so variable within a single ejaculate? Is the extensive transcription of non-coding DNA during spermiogenesis functional, or is it just an epiphenomenon? Finally, there are numerous translational applications of this technique as well, as a tool for validating diagnoses made from medical sequencing data, or for developing highly specific male contraceptives.

## Conclusions

Here we have reported a novel *in vivo* screening method to characterize the function of genes in the mammalian testis, and used it to validate 26 candidate genes as essential for spermatogenesis. It would have been impossible for our group to generate the same experimental evidence for these candidates using conventional single-gene mouse models. We have provided experimental protocols **(see File S1, File S2)** and an analysis pipeline **(see File S3)** to enable other interested groups to apply this screening technique to their own questions of interest.

## Methods

### Gene Selection

To design the pilot pool, we used data from the Mouse Genome Database (Eppig *et al*. 2015) from Jackson Labs (MGI) to create a list of genes that affect the male reproductive system when knocked out. We then used a list of genes that have been knocked out and not reported to cause any male reproductive defects to use as negative controls. For the predicted spermatogenesis gene pool, we picked the top 30 candidates from each of the mouse predicted infertility gene models (Ho *et al*. 2015) and filtered it to keep only the genes for which knockout mouse lines were not available based on the Jackson Labs MGI website. We then selected shRNAs against three of the known negative genes from the pilot pools together with two scrambled non-mammalian sequences to use as negative controls.

### shRNA Pool Preparation

We used RNAi from the MISSION^®^ TRC-Mm 1.5 and 2.0 (Mouse) obtained from Sigma-Aldrich **(see Table S1)** ordered as shRNA plasmids. The shRNA expression cassette from the plasmids was amplified from the plasmid pool by PCR and purified using AMPure XP beads **(see File S2)**. These purified amplicons were pooled and then used for injection into mouse testis and transfection into cell lines. An aliquot of this mixture was sequenced on MiSeq to determine initial shRNA pool composition.

### shRNA Validation

Over half (65/121) the shRNAs in the pilot pool and about a third of the shRNAs (48/145) in the predicted gene pool were reported by Sigma-Aldrich to be validated in various cell lines **(see Table S1)**. There was at least one validated shRNA for 72% (18/25) of the pilot pool genes and 45% (13/29) of the predicted pool genes. While not all the validated shRNAs were consistently significantly depleted in the two studies, many of them were, giving us confidence that the signal we observed was not caused by off-target effects.

### Mouse Testis Transfection

We performed the experiments using C57BL/6 mice generated in house between 28-32 days of age. All mice were maintained under pathogen-free conditions and all animal experiments were approved by Washington University’s Animal Studies Committee. Each mouse received bilateral intra-testicular DNA injections five times, spaced 3-4 days apart **(see File S1)**. Following the injections, the mice were allowed to recover until 20 days after the third injection, when testes were dissected. Genomic DNA from the whole testis was extracted using Qiagen DNeasy Blood and Tissue kit.

### Cell Line Transfection

N2a cells were maintained in Minimum Essential Media (MEM) supplemented with 10% Fetal Bovine Serum (FBS) until they reached 60% confluency in 6 well cell culture plates. Each well of cells was transfected with 2.5µg shRNA pool DNA using Lipofectamine^®^ 3000 (Life Technologies) following manufacturer’s instructions. The cells were then incubated at 37°C for 2 days, with daily media replacement, before the genomic DNA was extracted using the Qiagen DNeasy Blood and Tissue Kit.

### Immunofluorescence Assays

Testis tissues were fixed in 4% Paraformaldehyde diluted in PBS overnight. Following that, they were flushed in PBS 3 times for 1 hour each, followed by a flush in 30% ethanol for 1 hour, 50% ethanol for 1 hour and finally 70% ethanol for 1 hour. Processing of the tissue was carried out in serial flushing as follows: 70% ethanol for 5 min, 70% ethanol for 45 min, 80% ethanol for 45 min, 95% ethanol for 45 min, 95% ethanol for 45 min, 100% ethanol for 1 hour, 100% ethanol for 1 hour, Xylene for 1 hour, Xylene for 1 hour, Xylene for 1 hour, Paraffin for 1 hour, Paraffin for 75 min. Tissues were then embedded in paraffin blocks and sectioned at 5 micron thickness. Slides were boiled for 20 minutes in antibody retrieval solution (10mM Sodium Citrate, 0.05% Tween20, pH6.0) and stained with goat anti-GFP primary antibody (Abcam: ab5450) diluted 1:1000 in blocking solution (1X PBS, 0.2% Triton X, 5% Donkey serum) overnight at 4°C. A secondary antibody staining was performed using CF594 donkey anti-goat (Biotium: 20116) diluted 1:300 in blocking solution for 1.5 hours at room temperature. Hoeschst 33342 (Life Technologies: H3570) was diluted 1:500 in water and added to the slides for 5 minutes just before visualization.

### Illumina Sequencing Library Preparation

We used a custom protocol to amplify the shRNA sequences in the genomic DNA samples **(See File S2)**. This protocol used 2 rounds of PCR amplification to prepare the sequencing library instead of ligation followed by PCR amplification. We started with 2µg of genomic DNA to survey the genomes of enough cells in order to reduce the likelihood of dropout or PCR jackpotting; common artifacts when testing low numbers of cells.

Each biological sample had between three to five separate sequencing libraries prepared with different indices using different aliquots of genomic DNA to quantify technical noise. Sequencing libraries were then pooled and run in a lane of Illumina MiSeq 2x150bp to obtain an average of at least 3,000 reads per unique shRNA in the library.

### Statistical Analysis

Mapping of reads to shRNAs was done by aligning each read to the unique half of the hairpin sequence with no mismatches. A table of read counts for each shRNA was generated to determine significant enrichment/depletion. There was no significant difference in the counts between paired-end reads when they were mapped separately. We minimized technical noise by using the median value of technical replicates as the true count for each shRNA.

To determine significant depletion/enrichment of shRNAs, we used a custom R script **(S1 Script)**. We started by normalizing the shRNA count data to number of reads per shRNA per million reads in the sequencing library. We then calculated the log 2 fold enrichment of each shRNA in the testis relative to the initial DNA pool. The fold changes of different experiments using the same shRNA pool design were always normalized to the sequencing counts of the actual injected material. These fold changes were then merged to produce more biological replicates for a given shRNA pool design. Finally, we performed a Wilcoxon Rank Sum Test for each shRNAs’ fold enrichment across biological replicates against the fold enrichment of shRNAs against genes which are not known to affect spermatogenesis to calculate the likelihood that the shRNA was significantly depleted or enriched compared to the null. Any shRNA that had a p value smaller than the cutoff was determined to be significant. Due to the low transfection efficiency (1-3%) **(see Table S1)**, multiplicity of infection, a common issue in RNAi screens, was not a worry in the analysis.

### Network Analysis

We performed a functional pathway analysis of the genes targeted in the Predicted Genes screen and found four functional networks. We built networks based on co-expression, proteinprotein interactions, shared protein domains, and co-localization. We used Cytoscape (Shannon *et al*. 2003) with the GeneMANIA plugin (Montojo *et al*. 2010) with the default settings to visualize the functional network of the tested predicted genes **(see Figure S2)**.

## Data and reagent availability

The raw sequencing data supporting the conclusions of this article are available in the Short Read Archive under accession number SRP069249. Table S1 provides a detailed description of the design and clone name for every shRNA construct used in the study. Tables S3 and S4 provide normalized screening results for each shRNA construct. Table S5 provides the raw read counts for every shRNA construct from every experiment. The code that was used to analyze these raw reads counts and produce the results in this paper is provided in File S3.

## Author Contributions

NRYH, AU, YY, LM, and DC designed the initial composition and approach of the screening study. AU executed most of the mouse testes injections, while NRYH and YY performed some as well. NRYH prepared the material for injection and the samples for sequencing and quantification. NRYH and DC analyzed the data and prepared the manuscript.

## Acknowledgements

The authors thank Joseph Dougherty for providing materials and facilities for the N2a cell line experiments and Katinka Vigh-Conrad for assistance with preparation of two figures. We thank the Genome Technology Access Center in the Department of Genetics at Washington University School of Medicine for help with Illumina sequencing. The Center is partially supported by NCI Cancer Center Support Grant #P30 CA91842 to the Siteman Cancer Center and by ICTS/CTSA Grant# UL1 TR000448 from the National Center for Research Resources (NCRR), a component of the National Institutes of Health (NIH), and NIH Roadmap for Medical Research. This publication is solely the responsibility of the authors and does not necessarily represent the official view of NCRR or NIH.

## Supporting Information

**Figure S1. Overview of shRNA Library Prep**.

**Figure S2. Standard curve for Actin and shRNA qPCR primers** Slope for the actin curve is −3.797 with an R2 of 0.998, corresponding to 83.374% primer efficiency. Slope for the shRNA curve is −3.691 with an R2 of 0.996, corresponding to 86.614% primer efficiency.

**Figure S3. Functional networks for predicted genes** The black diamond nodes are genes that we tested in the predicted shRNA pool while gray circle nodes are genes predicted to be functionally related. Red gene names mean that the shRNA are annotated with GO terms that are related to sperm function, while black gene names have no such annotation. The four figures with identical nodes but different color lines indicate which GeneMANIA mouse network links the genes.

**Table S1. Summary of individual shRNA constructs used in this study**. Each row of the table corresponds to a single shRNA construct. For each construct, numerous details are provided regarding its design and validation. Two sheets are contained within the table, one for the Pilot Study and one for the Predicted Genes Study.

**Table S2. Comparison of Transfection Rates**. As described in the main manuscript, we performed a number of experiments to optimize the transfection of testicular germ cells. This table provides the raw data and calculations necessary to estimate the transfection rate for each experiment.

**Table S3. Summary Results of the Pilot Screen**. This table reports the summary results of the pilot screen in testis for each individual shRNA construct. The median effect size and p value for depletion are reported. “Effect” is calculate as the median log2 fold-change in the abundance of the construct in the testis library when compared to the input library. Thus a log2 fold-change < 0 indicates a depletion in the testis library compared to input, while a log 2 fold-change > 0 indicates an enrichment.

**Table S4. Summary Results of the Predicted Genes Screen**. This table reports the summary results of the predicted genes screen in testis for each individual shRNA construct. The format and interpretation are the same as for S3 Table.

**Table S.5 shRNA Read Counts**. The processed shRNA read counts from each experiment, derived from the raw sequencing data, are included within this table.

**File S1. Illustrated protocol for mouse testis injection**.

**File S2. Protocol for shRNA pool preparation and shRNA sequencing library preparation**.

**File S3. Code for analyzing shRNA count data**.

